# Experimental study of Young modulus of *Attacus atlas*, Vespa crabro, and *Libellula depressa* wings

**DOI:** 10.1101/390260

**Authors:** Michal Landowski, Zuzanna Kunicka-Kowalska, Krzysztof Sibilski

**Affiliations:** Gdansk University of Technology; Warsaw University of Technology

## Abstract

This paper describes a scientific research aimed at obtaining data for determining Young modulus of the wings of selected insects’ species. A small testing machine intended for three-point bending and equipped with instruments registering low forces was constructed for the needs of the experiment. The machine was used to perform numerous bending tests of wings of three species of insects (obtained from a breeding farm): *Attacus atlas, Vespa crabro, Libellula depressa* in various air-humidity conditions. Values of the force and displacement obtained in the course of the tests were used to calculate Young modulus. In order to do so, it was also necessary to obtain the moment of inertia of the wing cross-section. These values were measured on the basis of the images obtained with a SEM microscope. Obtained results were averaged and presented with a breakdown by air-humidity conditions. It was observed that Young modulus decreased with an increase of humidity; hence the calculations of the percentage decrease of this mechanical parameter were performed. Obtained results were compared with the observed structure which was also presented under light microscope. It transpired that the construction of a wing does not only influence the mechanical values but also it influences their susceptibility to the changes occurring in the environment. Thereby, differences between Lepidoptera and Hymenoptera insects were indicated also within the aspect discussed in this paper.

## Introduction

The paper contains a description of experiments performed in order to obtain the data necessary for calculating Young modulus of the wings of selected insect species. Research falling within the material science which have been performed so far focused mostly on the structure of insects wings instead of their strength parameters. Wings structures were examined, with a microscope as well, by scientists representing various disciplines, aiming at the description of the construction of a wing [1, 3, 4, 6, 9, 10, 12, 13, 14, 15, 17]. Some of these works attempt to re-create or mimic the wings, their deformations during flight or the frequency of natural vibrations, even with the application of the FEM method [13]. Others try to produce a material (a composite one, usually) reflecting the mechanical parameters and structure of the insect wings [1, 6] or even construct a Biomimetic Micro Aerial Vehicle (BMAV) that is a flying vehicle mimicking not only biological materials but also the way insects move [14]. A significant part of scientists focus solely on examining the construction of the wings of Lepidoptera insects. Microstructure of the particular scales and their fixings as well as the implication of such a construction are the subjects of numerous research and at the same time give hope of obtaining interesting results [3, 12, 17]. There are plans of taking the advantage of the knowledge on the influence of the scale structure on the colours of a wing. Among others, an example of Morpho butterfly was used to examine the method of reflecting white light of a scale in order to identify the method of creating a unique gloss on the wing. This knowledge can be used in manufacturing solar cells. There are also research projects which directly deal with the flexion of a wing and joints of its structures depending on the species of an insect [4]. As far as the research of insect wings is concerned, dragonflies and butterflies are most gladly used, followed by beetles. The latter constitute a very interesting case for flight mechanics, since one pair of wings evolved into elytron (wing covers) and in majority of species it is a fixed bearing surface, while the wings act as a propeller. [5]

Despite interesting and profound analyses, in the vast majority, these researches did not lead to obtaining reliable mechanical parameters. It is even harder because insect wings are natural composites and belong to anisotropic materials. Basically, the wings are constructed of veins which act as supporting frame and membrane spanning between these veins. The veins are placed in an irregular manner (neither longitudinal, nor transverse, at times almost radial – butterflies) [5]. What is more, each vein slightly differs in terms of construction. Individual features are also important. In case of large wings, it is possible to examine a specific section of a wing, while small wings need to be examined as a whole, without zonal division. This requires constructing a testing machine with equipment allowing bending specimens which are several hundred micrometers long and force sensors of a very small range. In their paper, Jiyu Sun and Bharat Bhushan [15] describe the construction of a dragonfly wing. They publish the photographs of the cross sections depending on the distance between the cross section and insect body along with the table containing mechanical properties: Young modulus and the hardness of various elements of wings. It is, however, the only such a vast source of data and it refers to one order of insects only. Additionally, the values are significantly diversified, depending on the wing element and measurement method. The research was performed with a nanoindenter which uses a diamond indenter to determine mechanical properties and defines the characteristics on the basis of nano-hardness measurements. Insect wings being a natural composite have a complex construction which is why determining the properties of a wing on the basis of the properties of the component materials without the influence of the reinforcement distribution and shape causes significant discrepancies with the real mechanical properties of the wings. Nevertheless, the data can be a used as a certain reference point and allow the verification of the correctness of performed experiments. Numerous literary sources contain information on the relationship between mechanical properties and condition of the insect wings. Young modulus value of a fresh membrane of an Allomyrina dichotoma hindwing ranges from 2.97 to 4.5 GPa. [16] Compared with the fresh hindwing, the Young modulus value of the dry membrane of a Allomyrina dichotoma hindwing is lower and varies over the area of the wing and ranges from 2.06 to 2.74 GPa. [11] These results suggest the necessity of selecting the air humidity conditions which will allow performing the research on mechanical properties compliant with the conditions of the insects habitats. Only the results of mechanical examinations performed in conditions which are similar to the real ones will allow using them in simulation of the wing movement and yield correct outcomes.

## Materials and methods

Due to the lack of appropriate measuring devices, in order to examine the insect wings a special testing machine adapted for biological material tests was designed and constructed. The appropriate selection of the components parameters was necessary because of low forces breaking the material and small sizes of the specimens. The machine consisted of two basic parts:

- Structural part: main frame made of aluminium profiles ensuring the appropriate stiffness, linear actuator and adjustable grip for the test of 3-point bending, made of S355 structural steel;
- Measurement part: force and displacement sensor with data acquisition system.

The initial design of the machine was made in Inventor software. The design which can be seen in fig. 01 did not assume the final selection of the elements; it rather aimed at specifying the dimensions and outline of the construction. Initially it was assumed that the displacement will be achieved with a gearbox transmission, however, an electrical linear actuator was used in the final version.

**Fig. 01.**
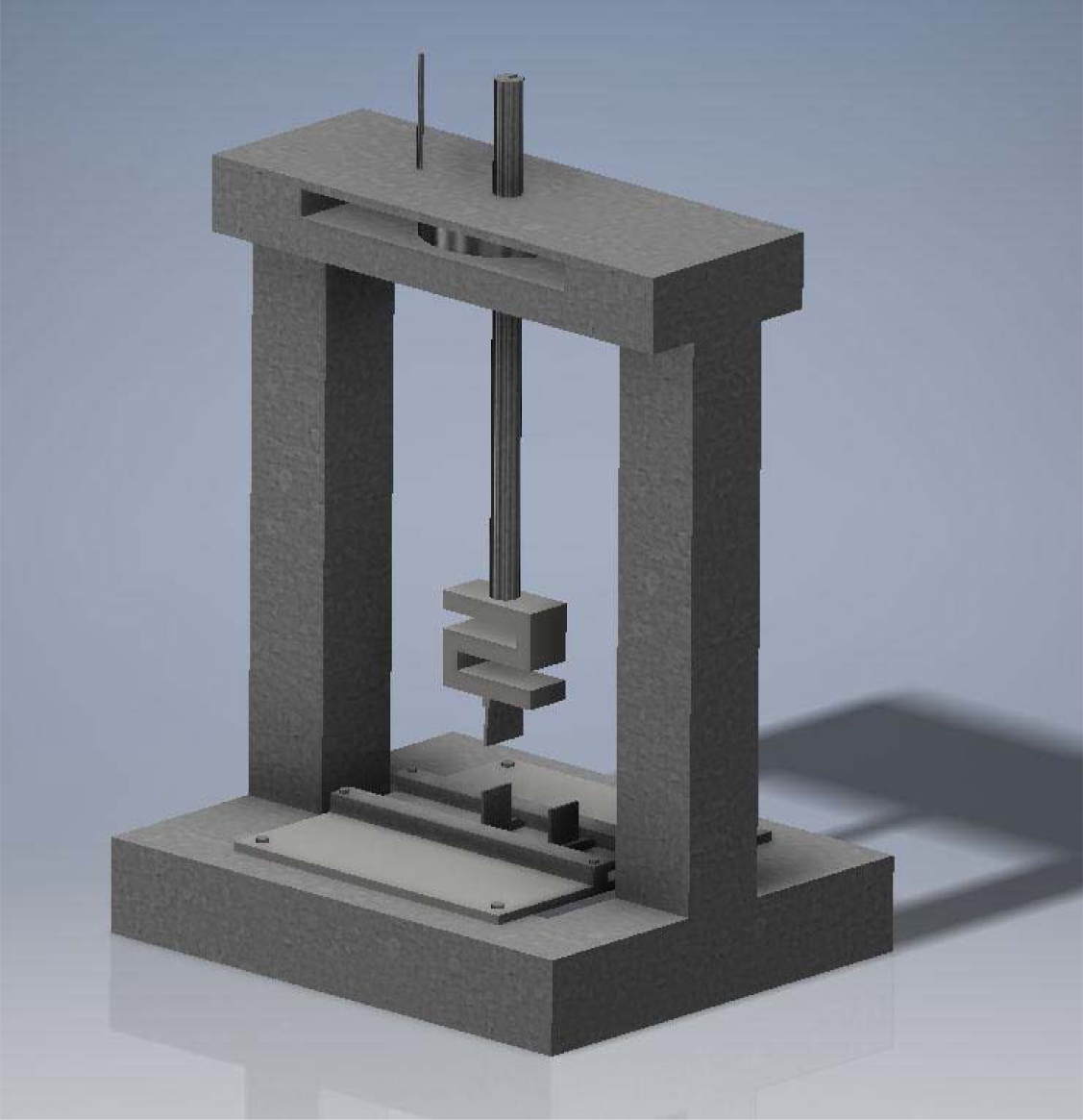
Initial concept and 3D outline of the machine

The main frame was made of 30 mm wide aluminium profiles featuring four T-slots, joined together with bolts and angle brackets to form the frame construction. Aluminium profiles used in construction ensured the stiffness of the structure due to their appropriate cross-section which translates into more precise results of the forces and displacement measurements. Using special profiles allowed fixing the elements with T-head bolts and arbitrary reconfiguration of the measurement system. Special cuts in the T-head of the bolts ensured self-locking and allowed the penetration of the oxide layer on the profile surface which ensures better dissipation of static electricity. Static electricity accumulated on the operator can significantly impede the measurements or even cause the damage of the measurement system. Electronic components in measurement system are powered by the current with 5V – 10V voltage, 100 V voltage is capable of causing a permanent damage of the components. In case of electrostatic discharge, the voltage exceeds 5kV - discharge of this type, at improper dissipation of the charge cause the majority of the damage in electronic components. The initial concept assumed transferring the progressive movement by a gearbox transmission joined with a trapezoid spindle translating the rotary motion of the motor into the progressive motion of the spindle. Such a concept assumed determining the pin stroke from the function of the revolutions of stepper motor. However, in the final version presented in figure 02, an electrical linear actuator DSZY4-12-50-100-IP65 with integrated DC motor powered by 12V DC, with 50:1 ratio internal gearbox, 100 mm stroke and maximum force of 2500 N was selected. Application of a linear actuator guaranteed the possibility of adjusting the displacement speed within the range of 0.25 – 5 mm/s for control voltage 0.5 to 12 V respectively, ensuring a more precise measurement.

**Fig. 02.**
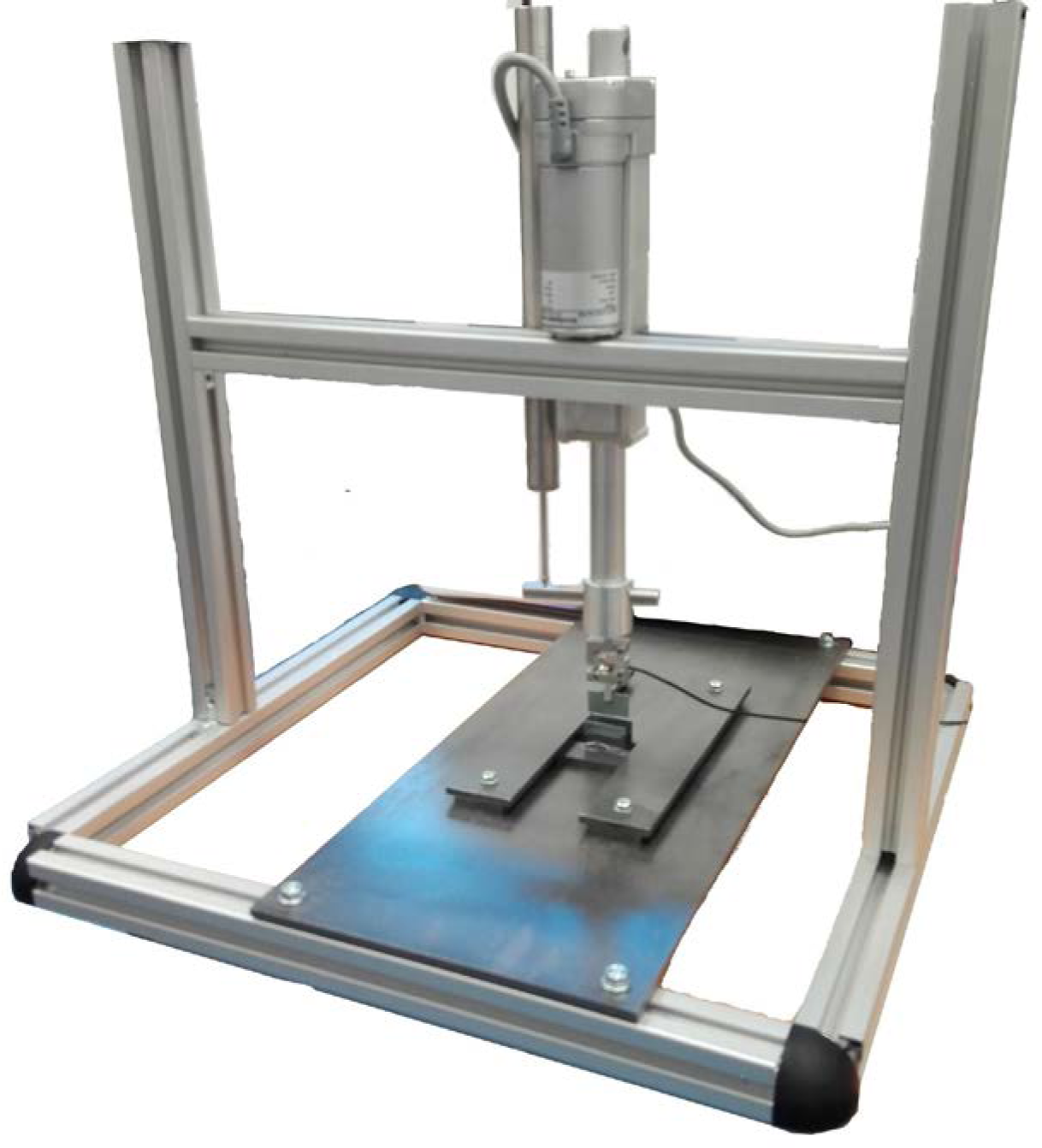
Measurement station

The advantage of a linear actuator consists in the possibility of avoiding abrupt displacement which is an obvious effect of using a stepper motor with a gearbox. The application of a linear actuator eliminated impact loads during bending (smooth displacement) which contributed to obtaining more reliable results. A metal grip for 3-point bending was fixed to the main frame. The grip was made of 5 mm thick plate with guides allowing smooth adjustment of the rests span. Particular elements were joined with bolts, allowing the modification and locking the setting of each element of the construction depending on the current needs.

Displacement sensor PELTRON PSz 50 with measurement range of 50 mm was fixed in parallel to the linear actuator. Displacement transducer is built on the basis of a differential transformer placed in a cylindrical housing, featuring a spring system allowing contact measurements. Transducer housing contains an electronic component with output signal of ±10V. The actuator was equipped with a tensometric S-shaped force sensor (for measuring the compressive and tensile forces) with a loading pin. Additionally, force sensor ZEPWN type CL14m, measurement range 2N powered with stabilized current of 5V and with 2mV/V sensitivity was selected. Measurement station presented in figure 03 features a 14-bit data acquisition system and separate stabilized power supplies in order to avoid the interference of signals from particular channels.

**Fig. 03.**
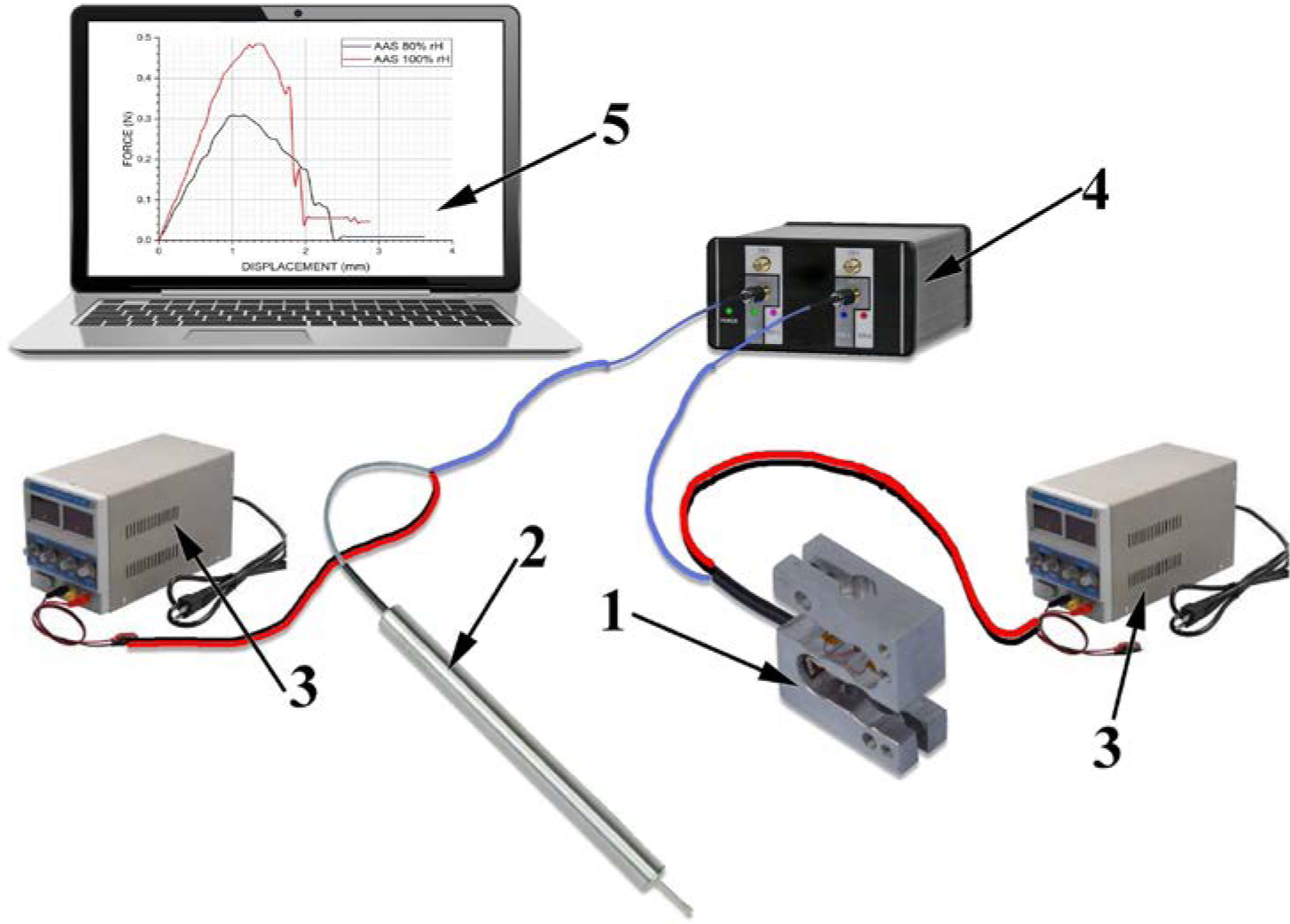
Schematic drawing of the measurement system. 1 –force sensor, 2 – displacement sensor, 3 – stabilized power source, 4 –signal amplifier, 5 – computer with data acquisition software

Three species of insects, one species of three various orders of winged insects: butterflies, Hymenoptera bees and dragonflies were selected for the examinations.:

- *Attacus atlas* (Fig. 04a) the largest representative of butterflies in the world. In the natural habitat it occurs in Ceylon, Malaysia, and China [2]. Specimens were obtained from a breeding farm.
- *Vespa crabro* (Fig. 04b) a representative of Hymenoptera bees, the largest representative of Vespidae in Poland. Its large wings facilitate the performance of the experiments.
- *Libellula depressa* (Fig. 04c) a common representative of *Odonata* (dragonflies) order widely spread in Poland.

**Fig. 04.**
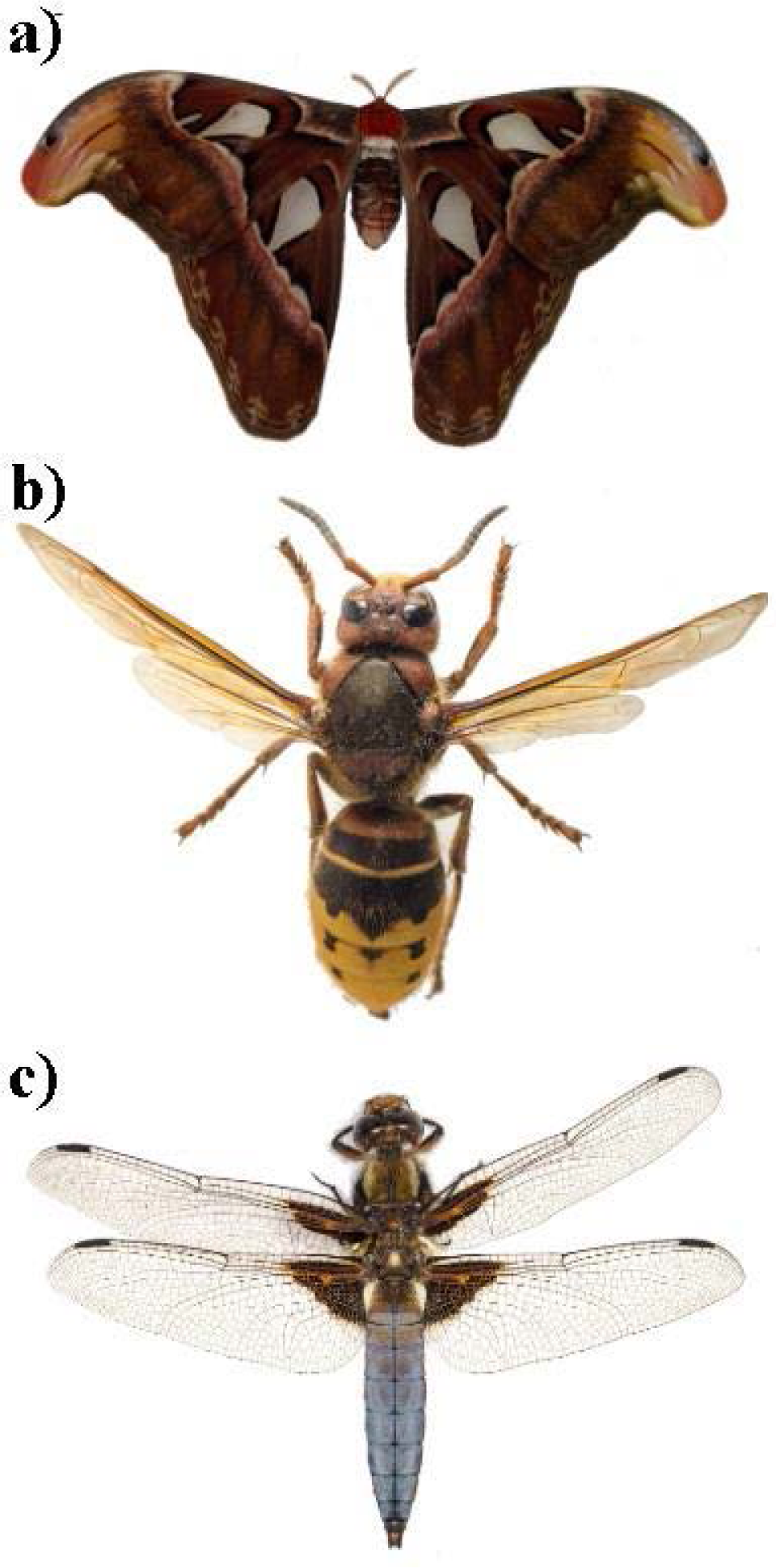
Reference photographs of the examined insects: a) *Attacus atlas*, b) *Vespa crabro*, c) *Libellula depressa*

All the specimens originated from the insects bought from a breeder. In the case of *Vespa crabro* and *Libellula depressa* deformation of the whole wing was measured. The hind wing of *Vespa crabro* was too small (too low strength) and the bending forces were not registered. In the case of *Attacus atlas* a section of the leading edge (Fig. 05a) and a section of the trailing edge (Fig. 05b) of a wing were examined. The examination of the butterfly wing membrane itself did not yield any effects: registered bending force equalled zero. The wing could have been, or even should have been divided into zones, since at such a large surface area, the mechanical properties exhibit significant differences depending on the location. Figure 05 shows examples of macro-photographs of the excised sections of wings which were the specimens under examination.

**Fig. 05.**
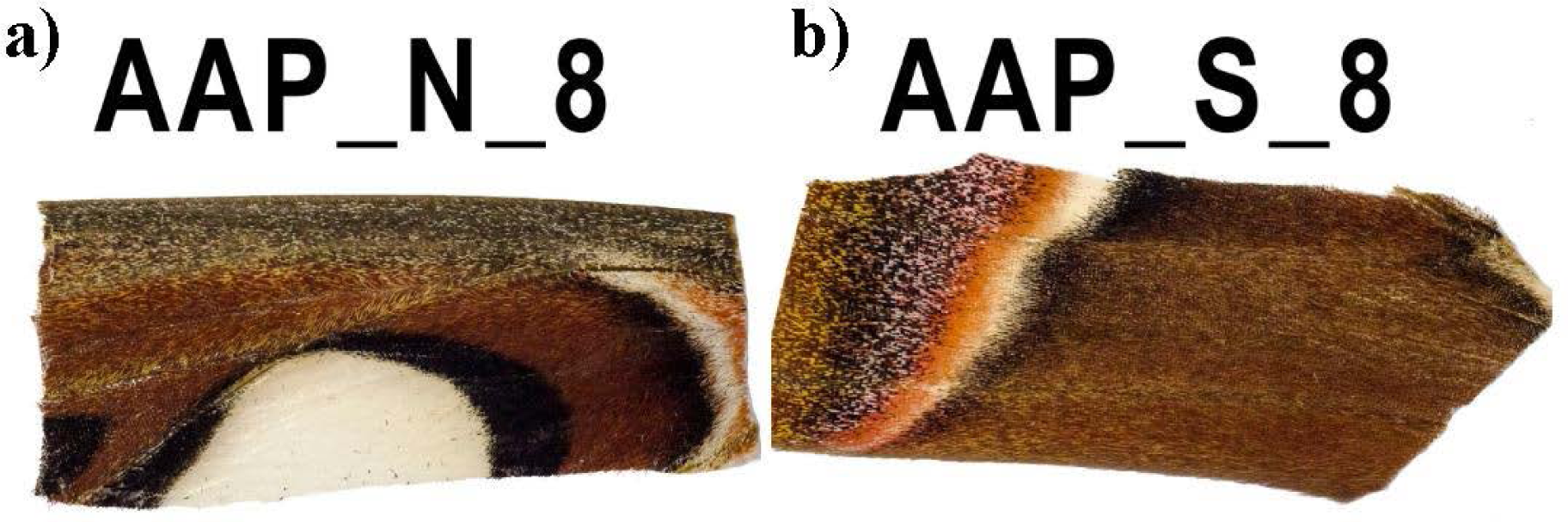
Excised sections of the forewings of *Attacus Atlas* a) leading edge (AAP_N_8), b) trailing edge (AAP_S_8)

Wing bending flexural stiffness was measured during the experiment by applying the force in the middle of the measurement section. The wing was being bent perpendicularly to its length. Examined specimens were placed on the rests freely, without any adhesive. The next stage consisted in slow lowering of the load pin to a level which was 2 mm above the specimen. Then, a stabilized power supply unit was used to set the voltage corresponding to the selected extension speed. In the next stage, the apparatus and data acquisition system were engaged and the voltage was supplied to the actuator. Force registration was initiated before the load pin contacted the wing and lasted until force value decreased after the break. The tested wing was unloaded by slow withdrawal of the load pin and it was put into safe storage with appropriate marking in order to examine the fracture with the SEM microscope. Power supply unit system was reset after each measurement and the correctness of the operation of force sensor was tested with three certified weighs: 1, 2 and 5 g.

The examination of the insects with more intricate or smaller wings, such as Tipula padulosa or smaller species of dragonflies was impossible since the breaking forces would be so low that they would not be correctly recorded or not recorded at all. Registration of such forces would require constructing a testing station isolated from the air movement (air would influence the measurements) and equipped with force sensor with range below 2 N. The experiment was performed in three various air humidity conditions in order to simultaneously specify their influence on the mechanical parameters of biological materials for whose the conditions of the operating environment have a significant meaning. The majority of specimens was conditioned at 80% air humidity and 25°C in Binder MKF115 climatic chamber for over 24 hours. Air humidity of 80 % reflects the condition of air prevailing in insects habitats in Poland. For comparison, the experiments included dragonfly wing specimens conditioned at approximately 100% and 30% air humidity. The humidity of air in the area where *Attacus atlas* is present reaches, approximately, 80 – 100% [8]. The examination at 30% air humidity had only theoretical character; it was not aimed at mimicking the natural conditions of the insects’ habitats. It was only performed in order to compare the mechanical values which biological materials could have in extreme conditions.

Bending tests were performed with two various settings of the lower rests: 5 and 10 mm span:

- For specimens of *Attacus atlas* and *Libellula depressa*: 10 mm,
- For *Vespa crabro* wing: 5 mm.

## Results

The measurements provided the relationship between the force and wing deflection. Figure 06 presents an example of the course of the force in the deflection function during the test of 3-point bending for specimens *Attacus atlas* leading edge in 80% rH and 100 rH conditioning. The graph exhibits a varied inclination of the initial, straight line section of the curve which indicates the differences in Young modulus depending on the humidity.

**Fig. 06.**
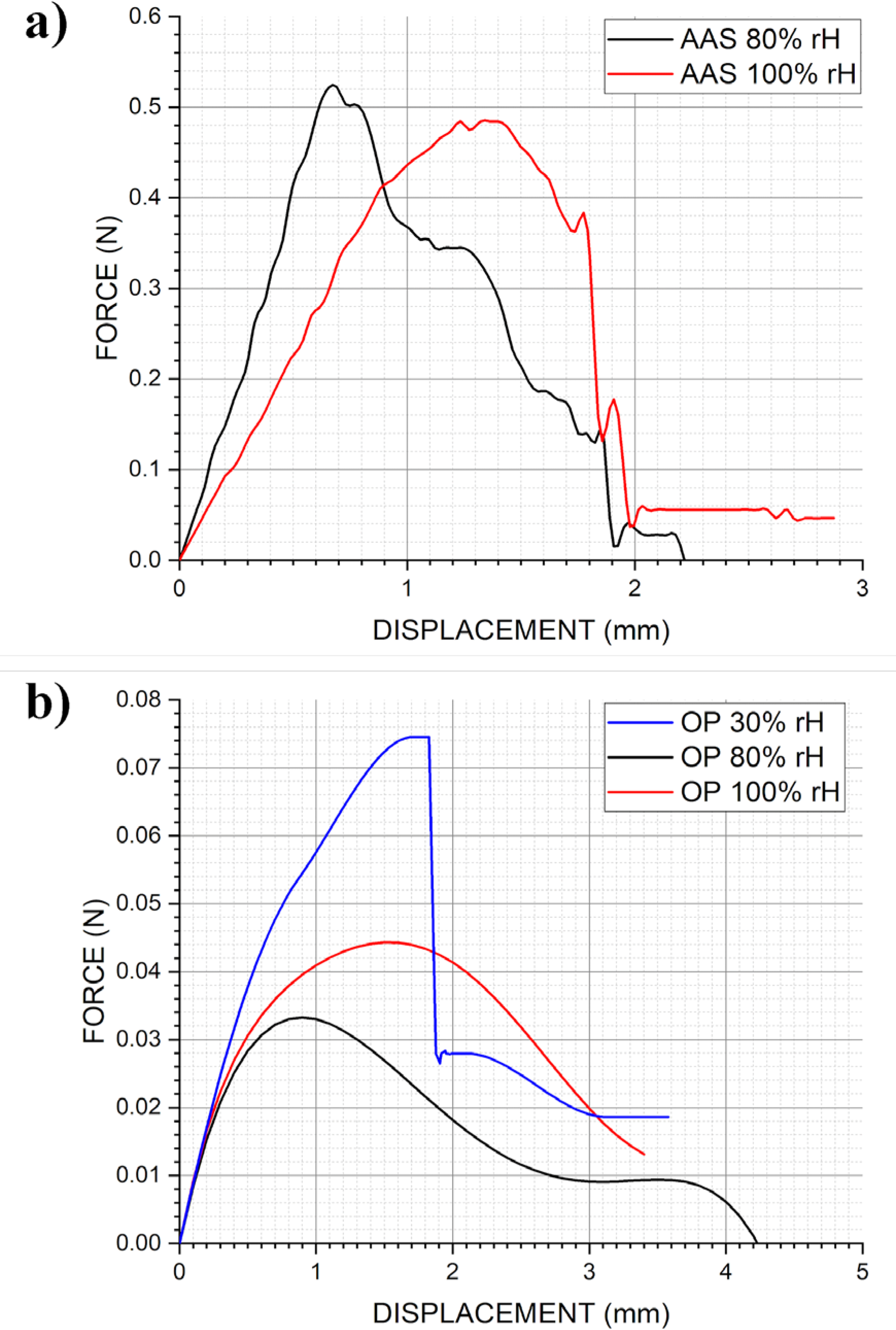
Example graphs of the force and displacement in various humidity conditions; a -*Attacus Atlas*, b - *Libellula depressa*

On the basis of the example graphs of the force and displacement it can be determined that the maximum forces obtained at bending 80% rH *Attacus atlas* specimen are lower than in case of specimens conditioned at 100% rH. Assuming similar cross-sections of the wings, this indicates a decrease of the flexural strength of wings at lower humidity. During the tests of a dry wing (30% rH) of *Libellula depressa*, a decrease of wing elasticity was noticed, there was a brittle fracture after maximum force was reached.

Fracture places of the wings and their surfaces were examined with Olympus BX52 light microscope. Figures below present the specific sections of the examined wings – their cross-sections and structure.

The structure of the wing surface of Lepidoptera insects differs significantly from other examined species. The difference is the most probable reason for varied susceptibility of the mechanical values to the changes in the humidity of the environment. Butterfly wings material is not as susceptible to the changes in the humidity. This is probably caused by the fact that the wings are covered with scales (Fig. 07a, b). The veins of the insects are made of chitin and their diameters are quite large in *Attacus* (Fig. 07c). In majority of cases the veins remain hollow, while smaller diameter veins are filled with protein. Due to this solution, it was possible to achieve high flexural strength with simultaneous minimization of mass. Very thin wing membrane whose thickness is in the order of 1 micrometer in *Vespa cabro* is covered with small bristles (Fig. 07d, e, f). This is probably connected with braking off a boundary layer of fluid during dynamic motion of a flapping flight. However, a precise description of the phenomenon would require profound examinations. Transverse cross-section (Fig. 07g) of a dragonfly wing shows that the wing is not a flat surface. Curves shown in this figure have significant meaning for the aerodynamics of flight. In spite of the considerable differences in sizes, the veins in Libellula *depressa* also remain hollow. Figure 07h presents a section of a wing with a stigma. This cell consists of two chitin walls and it remains hollow inside.

**Fig. 07.**
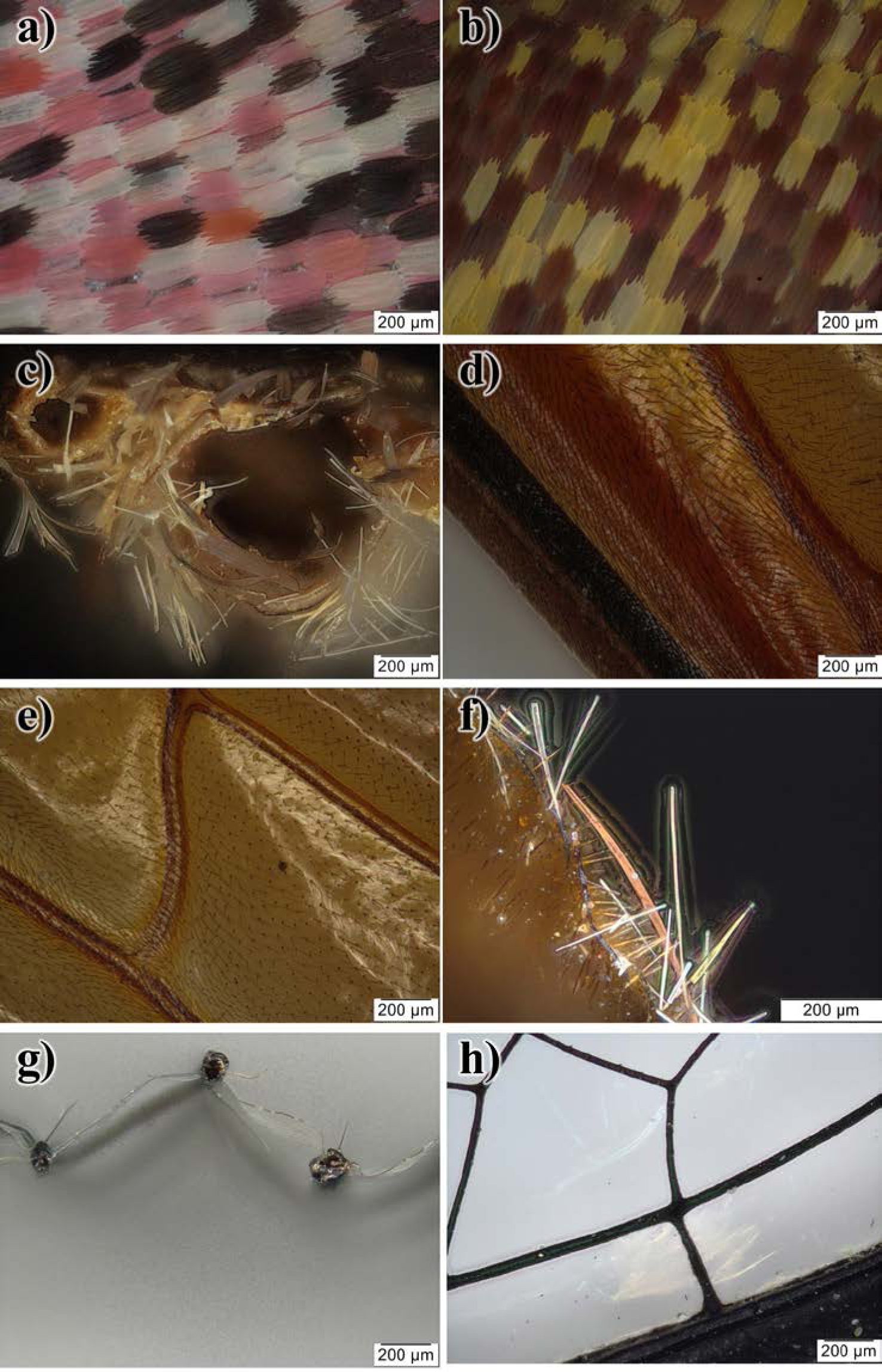
Microscopic photographs: a) *Attacus atlas* wing surface, section of a pink line, b) *Attacus atlas* wing surface, coloration change border, c) *Attacus atlas* cross-section of the leading edge, d) leading edge, e) *Vespa crabro* middle section of a wing, f) *Vespa crabro* cross-section of the membrane, g) *Libellula depressa* cross-section of the forewing, h) *Libellula depressa* surface of the forewing

General stiffness of a wing is the product of material stiffness (E, describing the stiffness of the wing material itself) and cross-section inertia moment (I, describing the stiffness generated by the geometry of the wing cross-section). In order to obtain Young modulus after bending tests, it was necessary to obtain:

- Flexion force - F,
- Rests span - L,
- Deflection value - s,
- Geometric inertia moment - I.

The relationship between deflection value and force was derived from the results of the experiments. Young modulus characterizes the stiffness of material within the range of elastic deformations following Hooke’s law. In order to calculate Young modulus it was necessary to consider only the linear relation from the first section of the graph. Exceeding the range of the deformation and stress proportionality is accompanied by non-reversible changes in the material – lasting deformation or rupture of the material continuity. The span of the rests was selected in relation to specific wings, set on the grip attached to the testing machine. The design of the measurement system allows smooth adjustment of the rests span; however two settings were sufficient for the examined specimens. They were adjusted to the size of the specimens in such a way as to ensure, on the one hand, that the specimen does not slide in between the rests (too big a distance) and, on the other hand, to avoid or minimize the shearing stresses occurring instead of bending stresses when the span between the rests is too small.

Inertia moment, being the measurement of a body inertia in rotary motion in relation to specific rotation axis could not have been calculated manually. Taking into consideration the irregularity of the shape, manual division into geometric figures of known inertia moment, finding the center of gravity and calculations considering Steiner theorem, would not only be burdened with a significant error, but also would be impossible to perform with the extent of precision allowing the result which would not interfere with further calculations. Thus, inertia moments were calculated in AutoCAD software on the basis of the microscopic photographs (Fig. 08) of the transverse cross-sections of wings at the breaking point. Photos were taken with a scanning electron microscope Jeol JSM-7800F. The outline of the cross-section was drawn manually, the envelopes of particular veins were combined into a single object. Next, with “physical parameters” command, geometric inertia moment was calculated automaticly.

**Fig. 08.**
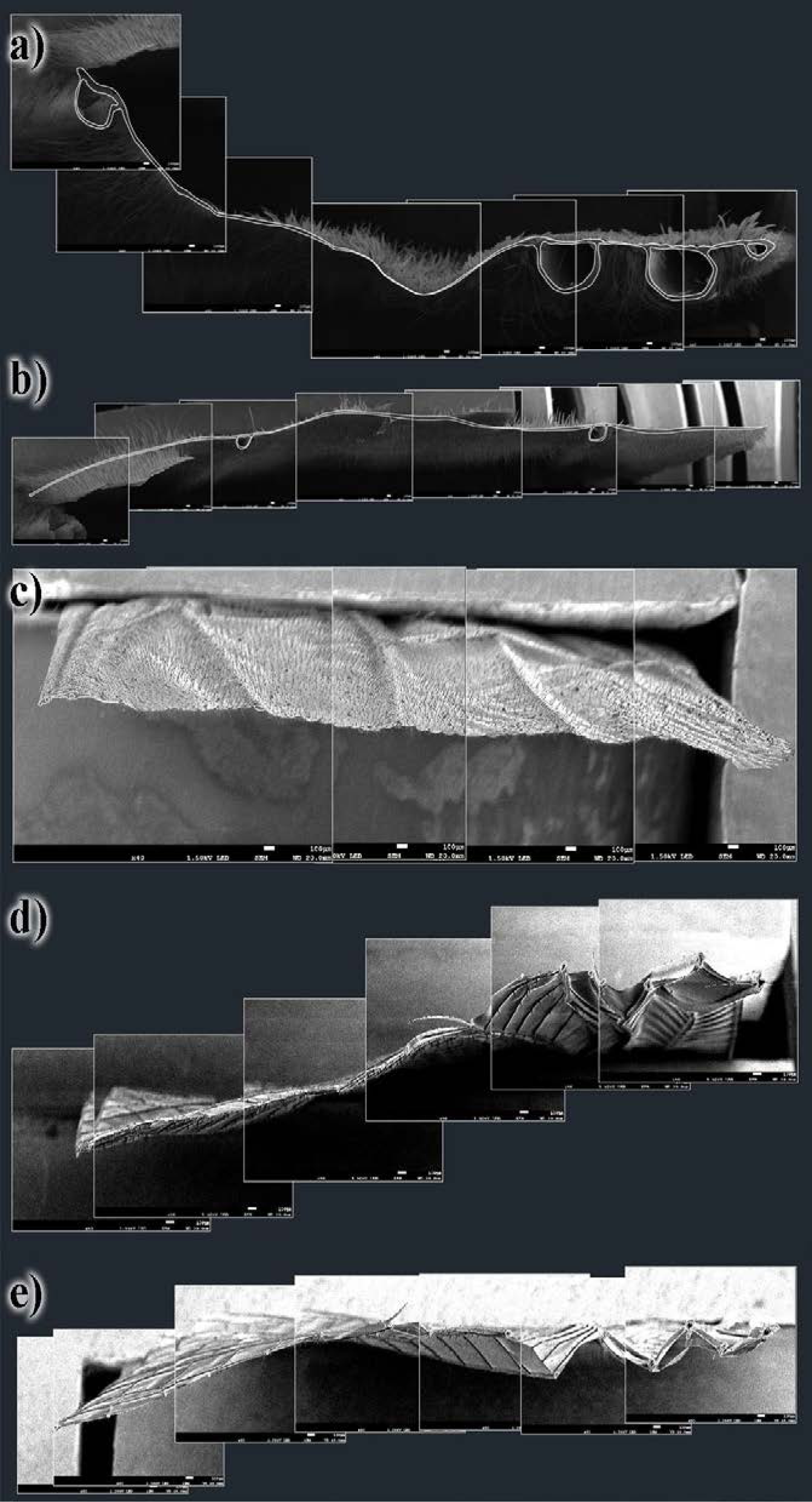
a) Cross-section of the specimen leading edge of *Attacus atlas*, b) Cross-section of the specimen leading edge of *Attacus atlas*, c) Cross-section of the specimen forewing of *Vespa crabro*, d) Cross-section of the specimen forewing of *Libellula depressa*, e) Cross-section of the specimen hindwing of *Libellula depressa*

Next, Young modulus was calculated with the formula below [7].

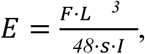

where:

*E* – Young modulus,

*F* – pressure force,

*L* – rests span,

*s* – deflection value,

*I* – geometric inertia moment.

The values of Young modulus expressed in GPa for three various levels of air humidity are presented below (Table 02).

**Tab. 01.**
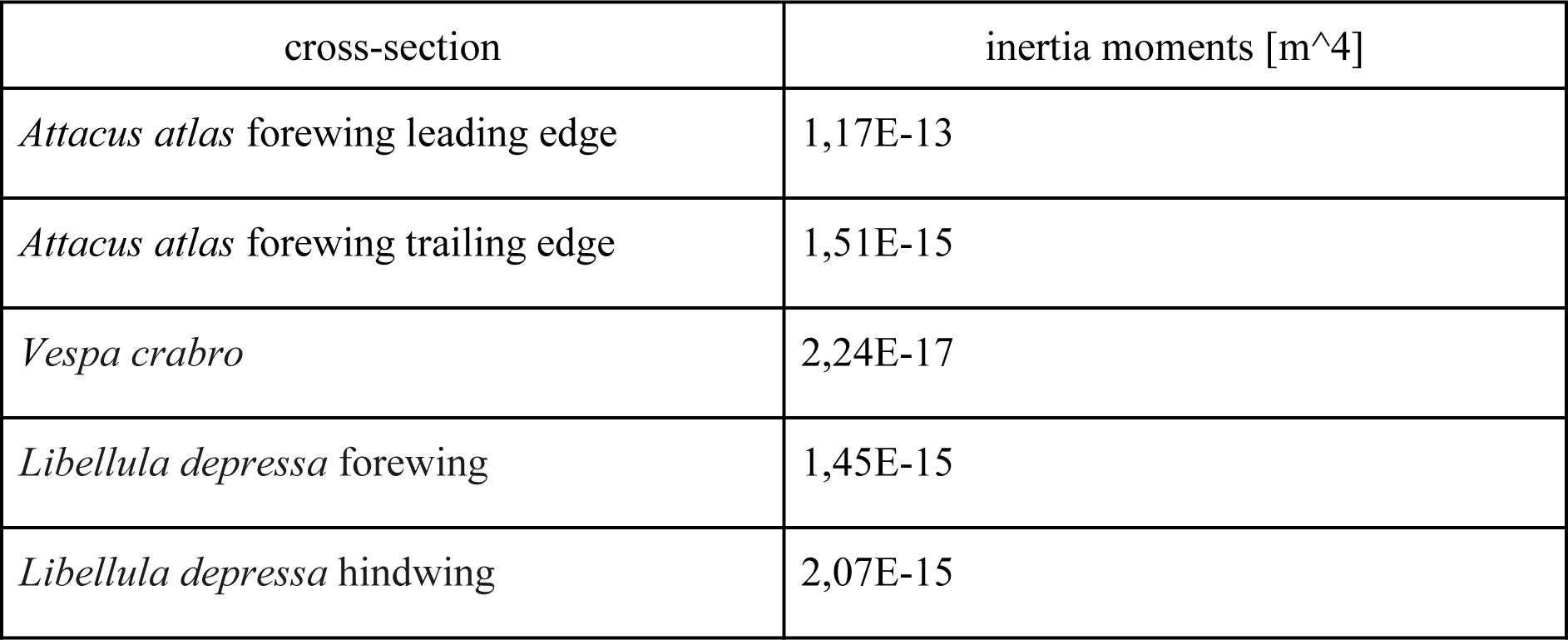
Inertia moments for cross-sections in Fig. 08

| cross-section | inertia moments [m<sup>4</sup>] |
| --- | --- |
| *Attacus atlas* forewing leading edge | 1,17E-13 |
| *Attacus atlas* forewing trailing edge | 1,51E-15 |
| *Vespa crabro* | 2,24E-17 |
| *Libellula depressa* forewing | 1,45E-15 |
| *Libellula depressa* hindwing | 2,07E-15 |

**Tab. 02.** Results of the calculations of Young modulus [GPa] and its percentage decrease

| insect | specimen of wing | 30% | 80% | 98% | percentage decrease |  |  |
| --- | --- | --- | --- | --- | --- | --- | --- |
|  |  |  |  |  | z 30 na 80 | z 80 na 98 | z 30 na 98 |
| *Attacus atlas* | forewing leading edge | - | 0,35 | 0,32 | - | 8,99 | - |
|  | forewing trailing edge | - | 7,45 | 6,54 | - | 12,20 | - |
| *Vespa crabro* | forewing | - | 13,88 | 6,72 | - | 51,55 | - |
| *Libellula depressa* | forewing | 1,81 | 1,65 | 0,86 | 8,98 | 47,64 | 52,34 |
|  | hindwing | 1,74 | 1,51 | - | 13,11 | - | - |

## Discussion

For comparison, Young modulus for examples of other natural materials falls within the following intervals [10]:

- balsa: 0,04-0,5 GPa,
- bamboo: 3-70 GPa,
- cotton: 8-35 GPa,
- apple skin: 0,06-0,08 GPa,
- leather: 0,007-0,08 GPa,
- egg shell: 11-40 GPa.

Natural materials are characterized by considerable differences in mechanical parameters. Also, in the case of Young modulus, lower and upper extreme yield a large interval. This results partially from the measurement errors, but mostly from the individual features of each specimen and each creature (or individual). This is the reason why it is necessary to average the data and results in order to perform further analyses.

During the described experiments, measurements yielded average values of Young modulus for various wings and at different humidity conditions. Subsequently, percentage decrease of Young modulus at the change of humidity was calculated. This way yielded a surprisingly repeatable result. When the value of air humidity changes from 80 to 100%, the decrease is approximately 10% for Lepidoptera insects and approximately 50% for the remaining wings specimens. Table 02 contains the precise data on the percentage decrease of Young modulus at the change of humidity level. The difference in the susceptibility to the change of humidity level should be detected in the structure of the wing. It transpires that covering the wings with scales significantly influences the wing mechanical properties and their change depending on the environmental conditions.

Essentially, the material of insects wings is exceptionally complex and its properties are unique and ontogenetic. Nevertheless, performed examinations indicated that it is possible to determine an approximate Young modulus if the wing is treated as a uniform material without determining the mechanical parameters of particular cells, veins or scales. It is a certain approximation, indeed, however it is necessary since the measurement of such small elements would be burdened with significant error and, what is more, would not have a considerable meaning for flight mechanism because the entire composite is engaged in the flight physics. Therefore, for the needs of this and the following research (e.g. on the flapping flight mechanism) it should be assumed that an insect wing is a uniform material of specific properties. Modelling of the particular parts of the structure would be pointless. Prospective clarification would be assuming the existence of the zones with various values of mechanical parameters. For example, in the case of large surface area of the wing of *Attacus atlas*, on the basis of the performed analyses it is possible to determine various values of Young modulus for the leading and trailing edges. However, it is impossible to examine the mechanical properties in a specific spot because, for instance, the examined specimen needs to have a specific length. For this purpose, it would be sensible to assume the existence of a certain distribution of Young modulus on the surface of the wing where the zones which were not measured directly would have values approximated on the basis of the measured ones. However, this will still be an approximation. Each attempt of recognition and mathematical description of nature needs to produce an approximation since we are only mimicking solutions which proved to be the best in the course of evolution.

